# Repeat sequences limit the effectiveness of LGT and favoured the evolution of meiotic sex in early eukaryotes

**DOI:** 10.1101/2022.03.22.485314

**Authors:** Marco Colnaghi, Nick Lane, Andrew Pomiankowski

## Abstract

The transition from prokaryotic lateral gene transfer to eukaryotic meiotic sex is poorly understood. Phylogenetic evidence suggests that it was tightly linked to eukaryogenesis, which involved an unprecedented rise in both genome size and the density of genetic repeats. Expansion of genome size raised the severity of Muller’s ratchet, whilst limiting the effectiveness of lateral gene transfer (LGT) at purging deleterious mutations. In principle, an increase in recombination length combined with higher rates of LGT could solve this problem. Here we show using a computational model that this solution fails in the presence of genetic repeats prevalent in early eukaryotes. The model demonstrates that dispersed repeat sequences allow ectopic recombination, which leads to the loss of genetic information and curtails the capacity of LGT to prevent mutation accumulation. Increasing recombination length in the presence of repeat sequences exacerbates the problem. Mutational decay can only be resisted with homology along extended sequences of DNA. We conclude that the transition to homologous pairing along linear chromosomes was a key innovation in meiotic sex, which was instrumental in the expansion of eukaryotic genomes and morphological complexity.

**Relevance Statement:** The origin of meiotic sex is a long-standing evolutionary enigma. This novel mechanism of reproduction replaced lateral gene transfer (LGT), the uptake and recombination of pieces of environmental DNA seen in bacteria and archaea. We link its origin to the expanded genome size and proliferation of genetic repeats found in early eukaryotes. Both factors led to high levels of mutation accumulation and gene loss under LGT which could not be retarded through increases in the rate of LGT or the length of DNA recombined. Meiotic sex with homologous pairing of long chromosome sized pieces of DNA promoted purifying selection and supressed ectopic recombination. It permitted the evolution of the expanded genome needed for the evolution of complex eukaryotic life.

## Introduction

The genes for meiosis are universal among eukaryotes, indicating that sex evolved before the divergence of the first eukaryotic clades (1, 2). It evolved from the molecular machinery for lateral gene transfer (LGT) which facilitates genetic exchange in archaea and bacteria (1, 3, 4). Prokaryotes possess homologues of the canonical molecular machinery for meiotic sex, including proteins of the *SMC* gene family of ATPases necessary for chromosome cohesion and condensation (5), as well as actin and tubulin, required for daughter cell separation and the movement of chromosomes (6). The *Rad51/Dcm1* gene family, which plays a central role in meiosis, also has high protein sequence similarity with *RecA*, responsible for homologous search and recombination in prokaryotes (7, 8). But why eukaryotes requisitioned this existing molecular machinery to evolve a completely new mechanism of reproduction, inheritance, and genetic exchange – meiotic sex – remains obscure.

Transformation is one of the major routes of genetic exchange via LGT in bacteria and involves the acquisition of environmental DNA (eDNA), followed by recombination into the host genome (8, 9). By allowing genetic exchange between lineages, transformation can restore genes that have been disrupted through mutation or deletion (10-12), counter the effects of genetic drift and reverse Muller’s ratchet (11, 13), and accelerate adaptation by reducing selective interference (14, 15). Previous modelling work has shown that the expansion of early eukaryote genome size was likely to have caused the failure of LGT (13). While LGT via transformation helps to purge deleterious mutations (11), this benefit rapidly wanes as genome size increases because of the difficulty of matching individual mutations with environmental DNA (13). LGT can resist mutation accumulation in larger genomes by combining more frequent recombination with increased recombination length, the mean length of DNA picked up from the environment and recombined into the host cell genome (13). But the distribution of recombination length in bacteria is skewed towards short eDNA sequences, with a median length that encompasses at most just few genes (16-18). In addition, bacteria typically cleave environmental DNA, shortening recombination length. While there are constraints on the rate of uptake and recombination through limited eDNA availability and sequence homology (12, 18), prokaryotes plainly did not follow the eukaryotic trajectory towards recombination across whole chromosomes.

After the endosymbiotic event which gave rise to the first eukaryotes, the archaeal host’s genome greatly expanded, enriched with genes of endosymbiotic origin and through gene duplication and divergence which enabled a range of novel functions (19, 20). This is estimated to have doubled gene number in LECA (21, 22). The extra energetic availability provided by the proto-mitochondrial endosymbiont released bioenergetic constraints over prokaryotic cell and genome size (23, 24). But this came with the cost of maintaining a larger genome (19, 21, 24, 25). Early eukaryote genome size expansion also reflected an increase in the density of repeat sequences, arising from gene duplication and the spread of mobile genetic elements (25, 26). Mobile retroelements of endosymbiotic origin are thought to have spread widely through the proto-eukaryote host genome, leading to a proliferation of self-splicing introns (27-29). These selfish elements are present in many bacterial species, almost always at low copy numbers (<10 per genome) (30), but likely increased in a more uninhibited manner, perhaps exploiting the nonhomologous end-joining mechanism of DNA repair found throughout eukaryotes (31). Novel intron density is thought to have reached a density comparable to that seen in modern eukaryote species (29).

The possible involvement of such repeat sequences has not been investigated in previous theoretical models of LGT or the transition to meiotic sex (10, 11, 13). In prokaryotes, high repeat density is associated with a high probability of ectopic recombination, increasing the rates of deletions, insertions, and other genomic rearrangements (32, 33). Recombination errors caused by the presence of repeat sequences introduce an additional cost to LGT, and potentially constrain the benefits of increased recombination length and LGT frequency. Here we investigate whether the sharp increase in repeat density in early eukaryotes could have forced them to abandon LGT in favour of meiosis. To investigate this hypothesis, we develop a computational model of mutation and selection in a population undergoing LGT via transformation in the presence of genetic repeats. The model highlights a trade-off between the benefits of LGT (greater genetic variance, enhancing purifying selection) and its cost (loss of genetic information through ectopic recombination). This leads to the view that the transition to meiotic sex was driven by the need for purifying selection in the expanding and repeat rich genomes of early eukaryotes, which could not be met by increases in recombination length or LGT rate.

## Results

### Muller’s ratchet

We use a Fisher-Wright process with non-overlapping generations to model the evolution of a population of *N* haploid individuals with a circular genome composed of *g* unique protein-coding genes, each of which can exist in either a wildtype or deleterious mutant state (**Figure S1; Table 1; Supplementary Methods**). The genome is interspersed at random intervals with a generic repeat sequence, at an initial density *ρ* per gene. Each generation, an individual has a probability *λ* of acquiring a fragment of eDNA of length *L*. Recombination requires matching of the terminal loci of the eDNA sequence and the host genome, either to a protein-coding gene or a repeat sequence. When there is multiple matching, one of the homologous sequences is randomly selected, with weights inversely proportional to the difference in length between the eDNA and possible matching host genomic sequences (**Supplementary Methods**). After LGT, deleterious mutations are randomly introduced at a rate *μ* per locus. The new generation is then generated by sampling with replacement from the old generation in proportion to individual reproductive fitness *w*_*m*_. The old generation dies and its DNA is released into the environment to constitute the eDNA pool for the next generation. At the end of each simulation, the severity of Muller’s ratchet is assessed through the rate of accumulation of mutations, deletions, and the total rate of gene loss.

**Table 1:**
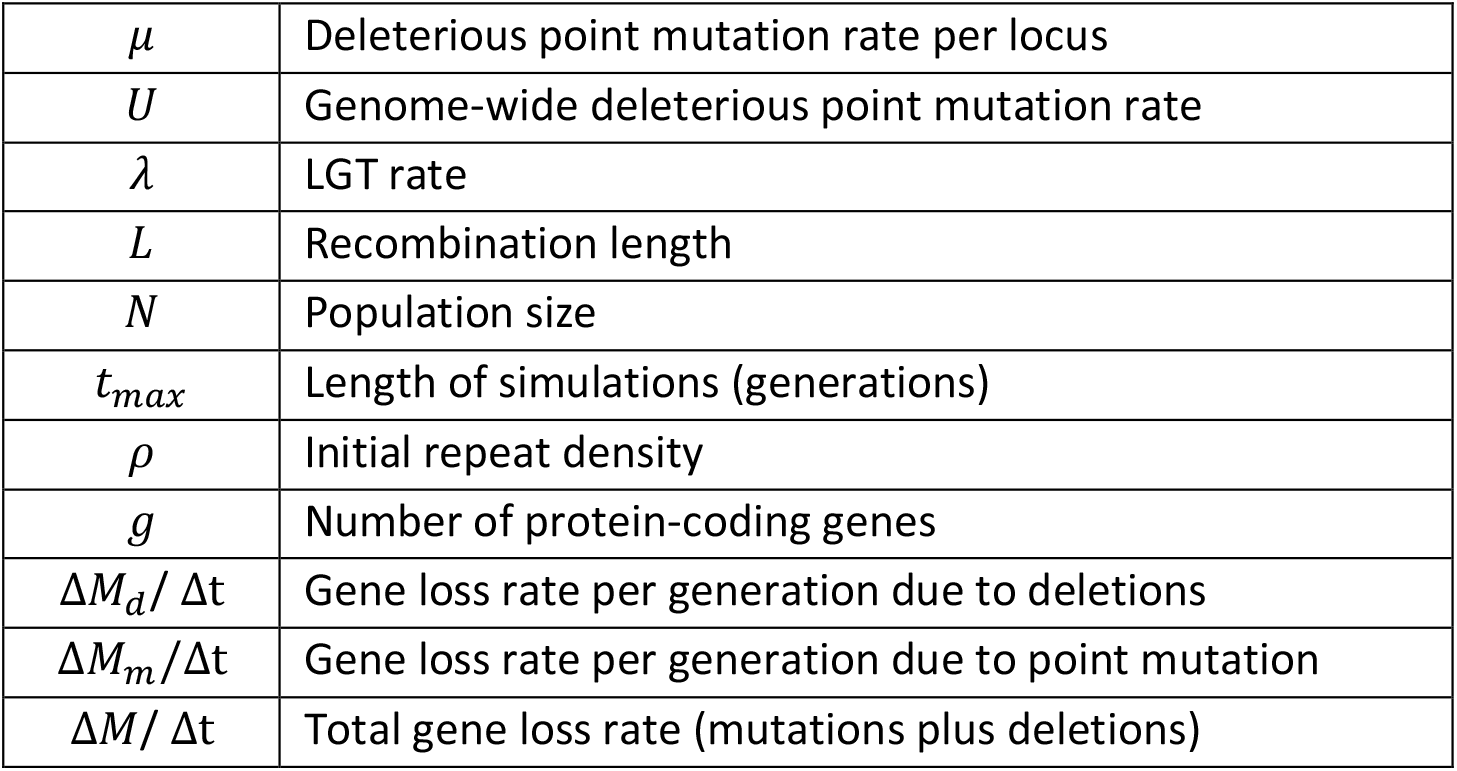
Model parameters and variables.

The mean mutation load of the whole population and the number of mutations in the least-loaded class (LLC) reflect the interplay between genetic drift and Muller’s ratchet (**Figure 1**). In a clonal population with no LGT (*λ* = 0), random fluctuations can cause the LLC to go extinct (a ‘click’ of the ratchet), determining an ever-increasing mutation load baseline (**Figure 1A**). In the absence of LGT, the fittest class cannot be restored, and the increase in mutation load is irreversible. The introduction of LGT (*λ* = 0.1) favours the elimination of mutations by increasing genetic variation and strengthening purifying selection, reducing the frequency at which the LLC is lost (**Figure 1B**). In addition, LGT permits the reversal of Muller’s ratchet and the reduction in mutation number in the LLC. But in the presence of a high repeat density (*ρ* = 0.1), the benefit of LGT comes at the price of deletions due to ectopic recombination, making LGT less obviously beneficial (**Figure 1C**).

**Figure 1.**
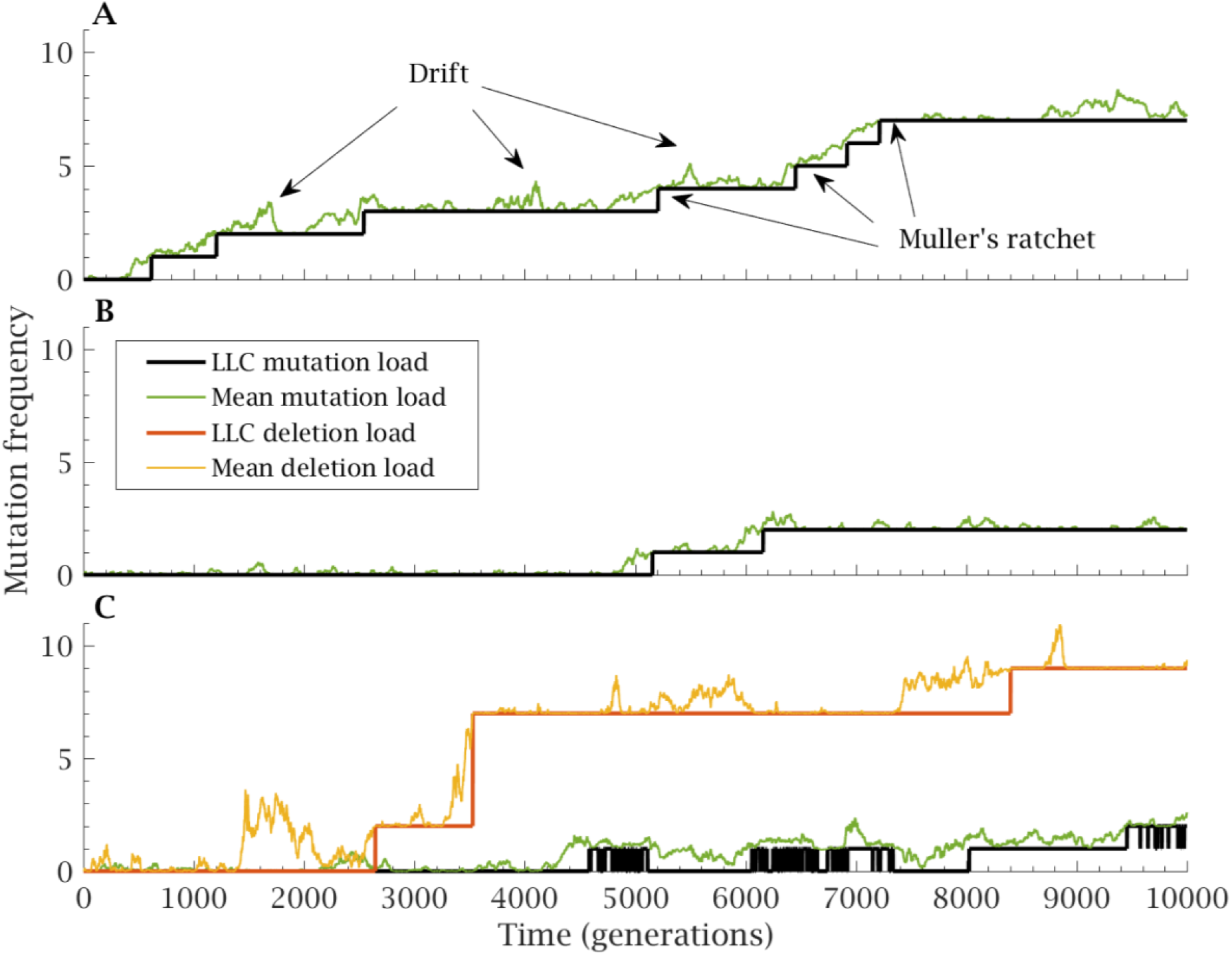
Impact of LGT on Muller’s ratchet. **(A)** Mean mutation load (green) and least loaded class (LLC) mutation load (black) of a repeats-free (*ρ* = 0), non-recombining population that does not undergo LGT (*λ* = 0), across *t*_*max*_ = 10,000 generations. Random fluctuations in allele frequencies due to genetic drift lead to irreversible increases of the LLC mutation load (Muller’s ratchet). **(B)** In a repeats-free (*ρ* = 0) population undergoing LGT (*λ* = 0.1, *L* = 5), recombination via LGT increases purifying selection, countering the ratchet. **(C)** In a population undergoing LGT (*λ* = 0.1, *L* = 5) in the presence of repeats (*ρ* = 0.1), LGT allows the ratchet to be reversed, reducing the LLC mutation load, but the presence of repeats leads to a high rate of deletion resulting in a high mean deletion load (yellow) and LLC deletion load (red). Other simulation parameters: *g* = 100, *N* = 2,500, *μ* = 3 × 10^−5^.

### LGT and repeats

Repeat density strongly influences the benefit of LGT. If repeat density is low (*ρ* = 10^−2^), increasing LGT (*λ*) is advantageous and reduces the total gene loss rate (**Figure 2A**). But as repeat density rises (*ρ* ≈ 0.5 × 10^−2^) this benefit is eroded, and higher levels of LGT provide little or no benefit (**Figure 2A**). At high levels of repeats (*ρ* ≥ 10^−1^), LGT is always detrimental and elevates total gene loss (**Figure 2A**). Splitting gene loss into its components, it becomes evident that the likelihood of ectopic recombination increases with higher density of repeats, leading to a sharp increase in gene loss through deletions (**Figure 2C**). In addition, high repeat density limits the effectiveness of LGT in purging deleterious mutations (**Figure 2D**). The net effect is that LGT ceases to have a beneficial effect beyond a threshold repeat density (**Figure 2B**).

**Figure 2.**
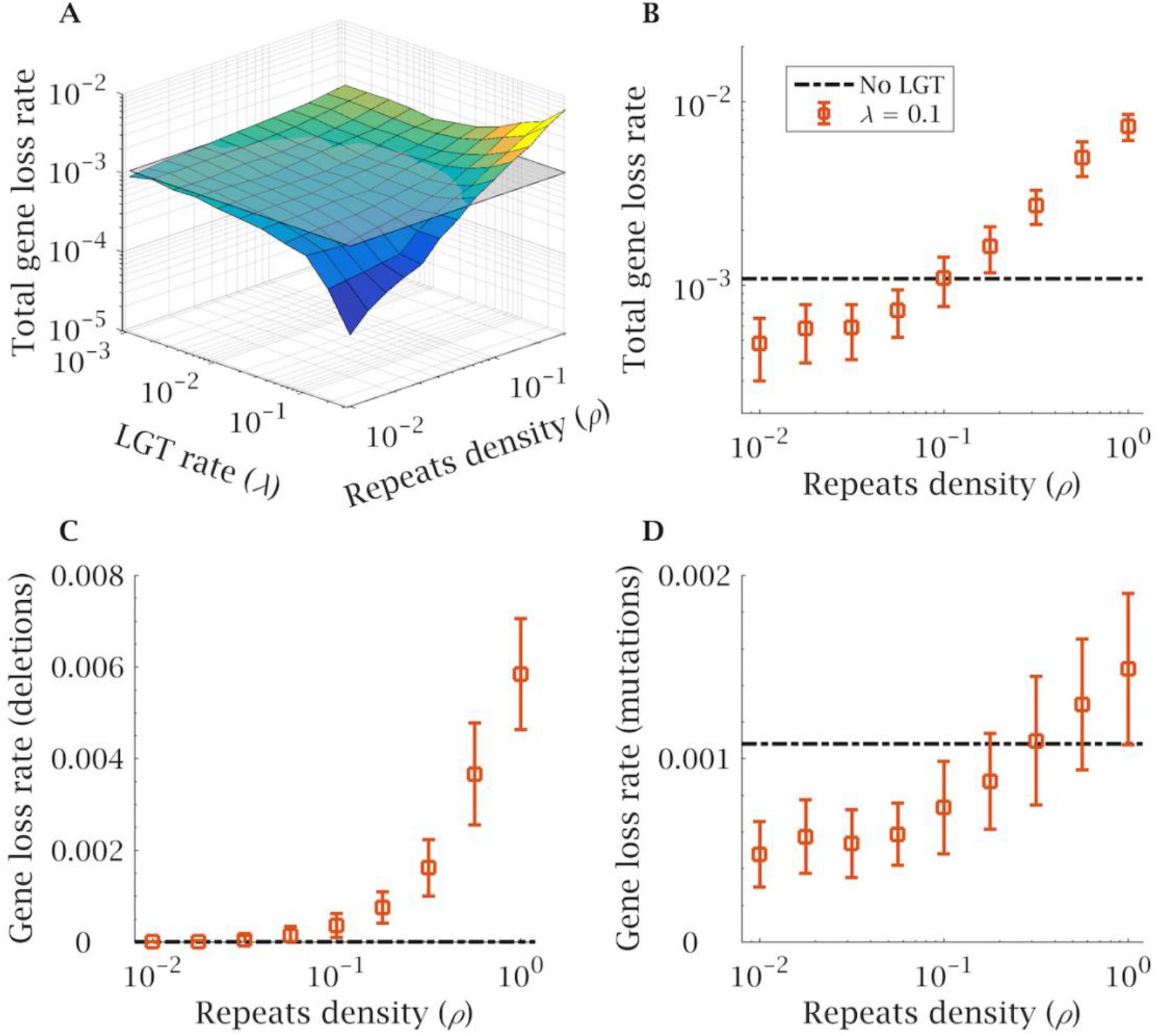
Repeat sequences cause the failure of LGT. **(A)** The total gene loss rate Δ*M*/Δ*t* (through both mutations and deletions) is shown for a range of initial repeat densities (*ρ*) and LGT rates (*λ*). For comparison, the grey plane shows the total gene loss rate in a repeat-free population not undergoing LGT (null model). Each data point is the average of 100 independent simulations. (**B**) The total gene loss rate (Δ*M*/Δ*t*) is due to (**C**) ectopic recombination leading to deletions (Δ*M*_*d*_/Δ*t*), and (**D**) recurrent deleterious mutations (Δ*M*_*m*_/Δ*t*), both of which increase with repeat density (*ρ*). Simulations in (**B-D**) were carried out with a high rate of LGT (*λ* = 0.1). The dotted line represents the null model of mutation accumulation in a non-recombining, repeats-free population. Error bars indicate the standard deviation over 100 independent simulations. Other simulation parameters: *g* = 100, *N* = 2,500, *μ* = 3 × 10^−5^, *t*_*max*_ = 5,000, *L* = 10.

The lower efficiency of LGT at removing deleterious mutations arises because repeats make homologous recombination less likely and ectopic recombination more likely. This can be seen by adding a requirement for full homology throughout the whole eDNA (i.e., not only at the terminal loci; see **Supplementary Methods**). This eliminates ectopic recombination, and mutation accumulation then closely follows the case without repeats, showing an exponential decline with LGT rate (**Figure 3A**). In contrast, with recombination based on end homology alone, there is only a monotonic decline in mutation accumulation as the rate of LGT increases (**Figure 3A**). This is because the presence of repeats, together with the build-up of deletions, raises the probability that one or both ends of the eDNA lack homology to any genomic sequence (i.e., if the matching sequence has been deleted from the genome, or if one end binds to a repeat sequence but the other end either lacks a homologous sequence or it too far away from it to recombine). Either scenario increases the probability that no recombination takes place (**Figure 3B**). These effects combine to reduce the rate of homologous recombination as repeat density increases (**Figure 3B**), constraining LGT’s ability to purge deleterious mutations.

**Figure 3.**
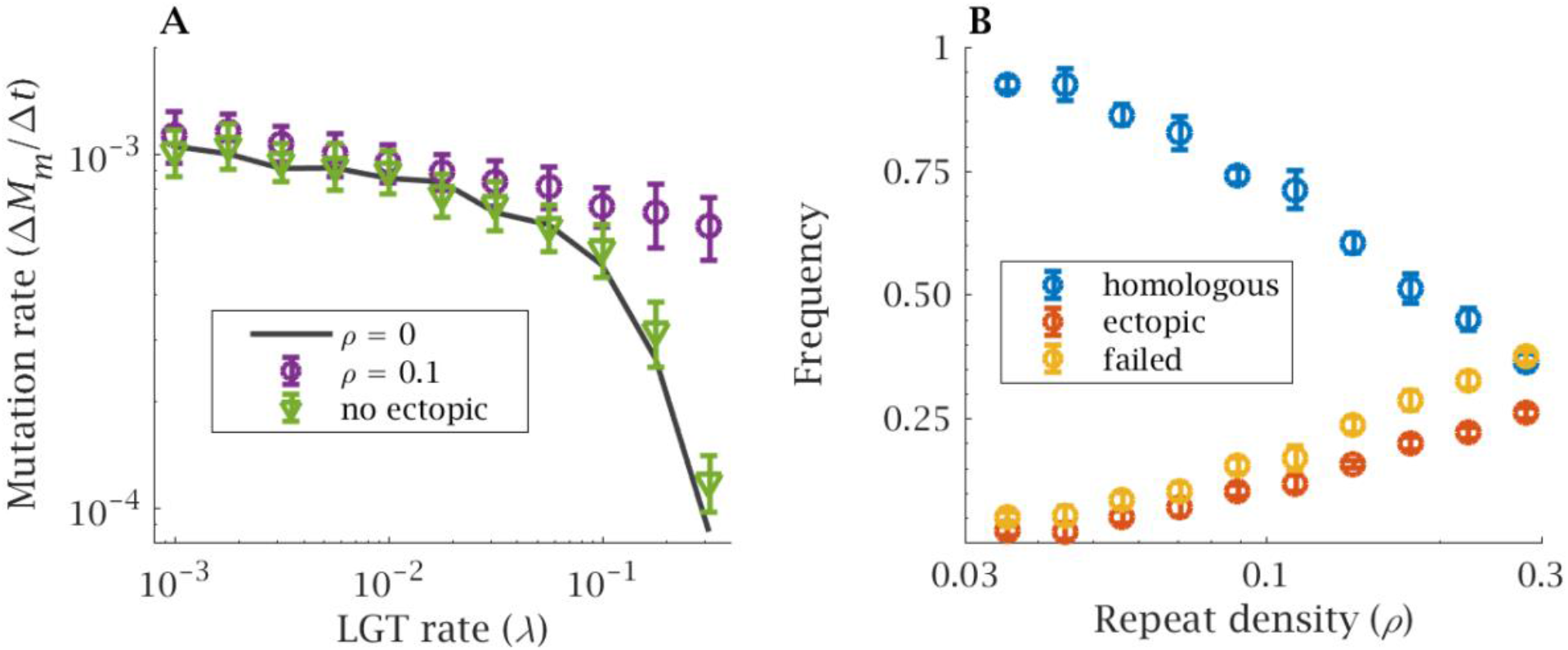
Ectopic recombination. (**A**) The rate of mutation accumulation is shown for the null model without repeats (*ρ* = 0, black line), a genome with high repeat density (*ρ* = 0.1, purple circles) and a genome with the same repeat content but where ectopic recombination is suppressed due to a requirement for homology throughout the eDNA (*ρ* = 0.1, green triangles, no ectopic). (**B**) The frequency of homologous (blue circles), ectopic (red circles) and failed recombination (yellow circles) as a function of repeat density. Rate and frequency were calculated over *t*_*max*_ = 5,000 generations. Error bars indicate the standard deviation over 100 independent simulations. Other parameters: *g* = 100, *N* = 2,500, *L* = 10, *μ* = 3 × 10^−5^ and in (**B**) *λ* = 0.1.

### Changes in genome size

The limitations of LGT are greater in large genomes (13). This is partly because the probability that eDNA matches a particular mutated sequence (reducing the mutation load) decreases with genome size. This effect can be overcome through increases in *L*, the size of eDNA selected for recombination (13). But that analysis neglected the effect of repeat sequences. If repeat density is low (*ρ* = 0.01), increases in *L* are favourable and help populations with large genome size resist the ratchet (compare *L* = 2 and *L* = 10; **Figure 4A**). However, with higher repeat density (*ρ* = 0.3), larger *L* is deleterious. It increases the total rate of gene loss and does nothing to stop the ratchet in large genomes (**Figure 4B**). This transition arises for two reasons. In genomes with few repeats, deletions through ectopic recombination are negligible and increasing *L* has only a very minor effect on their occurrence as genome size increases (**Figure S2A**). As almost all recombination events are homologous, increasing *L* is beneficial and facilitates the removal of deleterious mutations (**Figure S2B**). In contrast, when genomes are repeat rich (*ρ* = 0.3) the benefits of LGT are offset by an elevated rate of deletion (**Figure S2C**). This compromises the efficiency of eDNA repair of mutations, which becomes almost independent of *L* (**Figure S2D**).

**Figure 4.**
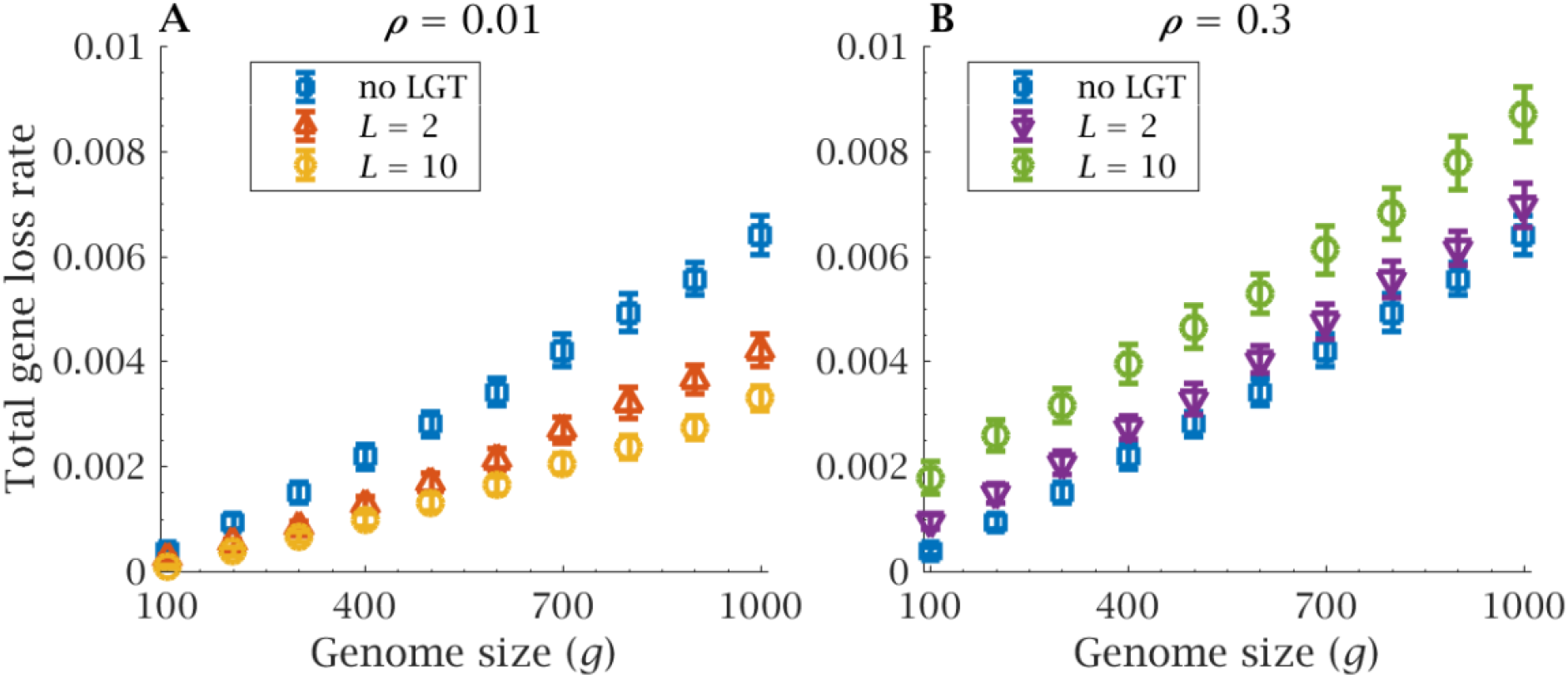
Genome size and recombination length. The rate of gene loss increases with genome size (*g*). (**A**) At low repeat density (*ρ* = 0.01), higher values of recombination length (*L*) minimise the rise in gene loss rate as genome size increase. (**B**) This benefit is reversed at high repeat density (*ρ* = 0.3), where higher *L* increases the total gene loss rate. Error bars show the standard deviation over 100 independent simulations. Note the null model (blue points) are identical in (**A**) and (**B**). Gene loss rate was calculated over *t*_*max*_ = 5,000 generations. Other parameters: *N* = 2,500, *μ* = 10^−5^ and *λ* = 0.1.

Unlike repeat-free populations (13), increasing eDNA length proportionally to genome size provides little or no benefit in the presence of repeats. A large genome (*g* = 1,000) cannot be sustained by LGT above a critical repeat density, as this engenders a rate of total gene loss comparable to that of a non-recombining population (**Figure 5A**, red line). The way out of this dilemma is to require sequence homology throughout the eDNA. This ensures that recombination is homologous and allows a reduction of the mutation load with large *L* without incurring the associated increase in gene deletions. As a consequence, recombination is able to lower the total gene loss rate in large (*g* = 1,000) genomes even in the presence of a high density of repeated sequences (**Figure 5A**, yellow line).

**Figure 5.**
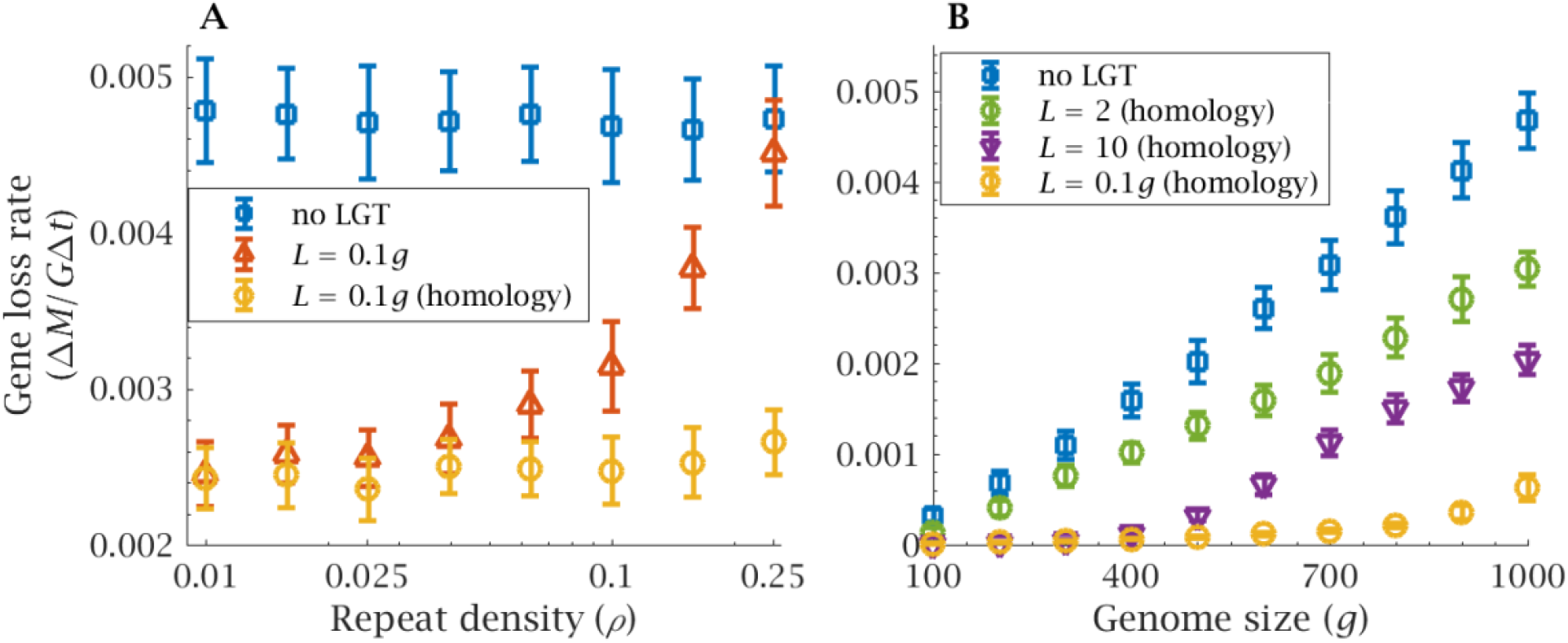
Advantage of homologous recombination. (**A**) Total gene loss rate as a function of repeat density (*ρ*). The blue squares correspond to in the null model of a non-recombining population, where the total gene loss rate is independent from repeat density. This is compared to LGT of long eDNA sequences (*L* = 0.1*g*, red triangles) and recombination with full sequence homology throughout the eDNA (yellow circles) with the same recombination length. (**B**) The impact of recombination length (*L*) on total gene loss as a function of genome size (*g*) in a repeats-rich genome (*ρ* = 0.3) under the requirement for full sequence homology throughout the eDNA. To limit gene loss in large genomes, recombination length must increase proportionally to genome size (*L* = 0.1*g*, yellow circles). Error bars show the standard deviation over 100 independent simulations. Gene loss rate was calculated over *t*_*max*_ = 5,000 generations. Other parameters: *N* = 2,500, *μ* = 10^−5^; in (**A**) *λ* = 0.1, *g* = 1,000 and (**B**) *λ* = 1, *ρ* = 0.3.

Recombination lengths proportional to genome size (*L* = 0.1*g*), homologous recombination across the entire length of the eDNA, and a high LGT rate (*λ* = 1) are all needed to prevent a sharp increase in total gene loss associated with a large genome size (**Figure 5B**, yellow line).

## Discussion

Asexual organisms are often portrayed as destined to accumulate mutations via Muller’s ratchet, on an inevitable decline to extinction through mutational meltdown (34-36). This view emanates from a eukaryotic perspective seeking to explain the maintenance of sexual reproduction in the face of the two-fold cost of sex and other costs relating to meiosis, finding a mate and cell fusion (37-39). Prokaryotes, both bacteria and archaea, lack meiotic sex and typically reproduce through asexual division, but they nonetheless have a number of mechanisms for achieving genetic recombination (12, 40-42). In particular, lateral gene transfer (LGT) through transformation allows competent cells to pick-up eDNA released from related lineages and to recombine it into their genome. In agreement with previous studies (10, 11, 13), our modelling shows that LGT generates genetic variation, strengthens purifying selection, and reduces the rate of mutation accumulation, performing a similar function to meiotic sex in eukaryotes [standard REFS]. This leads to a simple question: why did the first eukaryotes abandon LGT and replace it with meiotic sex?

Our analysis shows that the benefits of LGT are curtailed by the presence of genetic repeats (**Figure 2A**). Repeat sequences enable ectopic recombination, causing gene loss through deletions proportional to their density in the host genome (**Figure 2C**). Repeats also make LGT less effective at purging deleterious mutations (**Figure 2D**) by reducing the rate of correct homologous recombination (**Figure 3**). In addition, the presence of deletions and mutations segregating at the same time increases selective interference (43, 44), lowering the effectiveness of selection (**Figure 3**). A higher deletion rate, lower homologous recombination rate and weaker selection amplify the loss of genetic information as repeat density increases (**Figures 2-3**).

The acquisition of new genes through endosymbiotic gene transfer (including mobile self-splicing introns), plus duplication and divergence, led to massive genome expansion in the evolution of early eukaryotes (21, 25, 26). This made the first eukaryotes more vulnerable to the accumulation of mutations caused by Muller’s ratchet. As genome size (*g*) rises, the homologous recombination rate *per locus* falls, simply because the probability that an eDNA piece matches to a particular locus is inversely proportional to genome size (13). A solution to this is to increase the recombination length (**Figure 4A**). Picking up larger pieces of eDNA (larger *L*) increases the recombination rate *per locus* and thereby facilitates the elimination of deleterious mutations (13). But this potential solution is compromised by the presence of genetic repeats. If repeats are common, larger recombination length (*L*) is associated with higher, rather than lower, loss of genetic information (**Figure 4B**). A higher recombination length elevates the rate of ectopic recombination between dispersed repeated sequences, resulting in a greater rate of gene deletions. Similar disadvantages have been reported in extant prokaryotes where higher repeat density is associated with greater genomic instability, increasing deletions, inversions and other genomic rearrangements (32, 33). All of these issues are amplified as genome size increases, and beyond a threshold in repeat density, LGT brings no benefit to a large genome even if the recombination length scales with genome size (**Figure 5A**).

In order to support an expanded genome rich in repeat sequences, the first eukaryotes had to abandon LGT for meiotic sex. The transition to cell fusion and the requirement for homologous pairing across whole chromosomes was the only way they could retain the benefits of recombination without losing genetic information through LGT in the presence of repeats. We simulated meiotic sex in our LGT model by adding a requirement for homology throughout the eDNA as well as homology at the ends. Homology matching eliminates gene deletion through ectopic recombination and allows mutation accumulation to be resisted even as repeat density increases (**Figure 5A**). Considerable expansion of genome size is now permissible without catastrophic loss of genetic information through Muller’s ratchet, provided that recombination length (*L*) scales with genome size (**Figure 5B**). This is equivalent to homologous recombination across aligned chromosomes as seen in meiosis, though this neglects meiotic exchange in meiosis rather than replacement in LGT.

For simplicity, our model considers only one type of repeat sequence. In reality there will have been numerous repeats at different densities. Frequent gene duplications in early eukaryotes contributed to the increase in repeat density, and it is estimated that the average copy number per gene in LECA was around 1.8 (25). Another source of repeat sequence in prokaryotic genomes is transposable elements (TEs) and other selfish genetic elements. TEs can promote their own spread and reduce host fitness in other ways, through gene function disruption or gene inactivation (45, 46), and are thought to play a major role in the streamlining of prokaryotic genomes (47). The density of mobile intron-derived sequences in ancestral eukaryotic genomes is estimated to be as high as 80%, making the choice of *ρ* = 0.1 in most of our simulations a conservative one (48). As the focus of this study is on recombination and genetic information loss, we did not explicitly model the population dynamics of repeats, but evaluated the effect of variation in their initial density. Future studies should consider a diversity of repeats and include fitness penalties at the individual level associated with their movement and density, as well as the expansion of selfish genetic elements through replication within genomes. In itself this raises interesting questions about the distribution of repeats in extant prokaryotes – for instance, why the distribution of recombination length in prokaryotes is skewed towards shorter sequences (18), why gram-positive bacteria cleave eDNA sequences before recombination (49) and why the number of transposable and other mobile elements in prokaryote genomes is so tightly constrained (30, 50).

A further possibility not covered by our modelling are beneficial ectopic recombination events, in particular, those that contribute to the acquisition of novel genes. LGT via plasmids is the main source of acquisition of accessory genes from distant lineages (51-53) and has been shown to provide adaptive benefits (54). In contrast, transformation is mainly limited to sequences from closely related lineages, requiring a high degree of sequence homology and so less likely to import foreign genes across large taxonomic distances (55-58). As such, the main advantage of transformation is believed to be maintaining local adaptation rather than import of novel functions (9). In our model, gene loading can occur via the acquisition of genes lost through deletion. However, in the presence of a high repeat density, this effect is negligible compared to the loss of genes via deletions. It seems unlikely that repeats are retained in order to enhance gene turn-over from the pan-genome, but a proper analysis of this question would require a different modelling approach (59, 60).

Wilkins and Holliday (61) suggested that meiosis could arise from mitosis in a single evolutionary step, the evolution of homologous pairing during prophase. Our analysis here indicates that homologous pairing could have arisen because of the need to evade the deleterious effect of pervasive genomic repeats in the ancestral eukaryotic genome. Whilst our analysis demonstrates the selective advantage of meiotic sex, we do not explicitly address the multi-faceted question of its evolution. After the acquisition of mitochondrial symbionts allowed eukaryotic genome size expansion (23, 24), what promoted cell fusion and the reciprocal exchange across the whole genome, when did pre-meiotic doubling and a two-step meiosis evolve (61, 62), what selective forces imposed the haploid-diploid system of reduction division and why was meiosis the solution for an expanded archaeal nuclear genome (63, 64), while the endosymbiotic bacterial genome shrank almost to oblivion, lost capacity for LGT and became a multiploid, asexual, uniparentally transmitted mitochondrial genome (65-67)? All those steps were crucial for the survival and evolution of early eukaryotes. Without them, complex life as we know it could not have survived its inception.

Nonetheless, our work here shows why early eukaryotes had to take up whole chromosome-sized pieces of DNA and align them along their full length, rather than simply end-matching, clarifying the first necessary step from LGT towards meiosis.

## Acknowledgements

This work was supported by funding from the Engineering and Physical Sciences Research Council (EP/F500351/1, EP/I017909/1) and Natural Environment Research Council (NE/R010579/1) to AP, the Biotechnology and Biological Sciences Research Council (BB/S003681/1) and bgc3 to NL, and a joint grant to AP and NL from the Biotechnology and Biological Sciences Research Council (BB/V003542/1).

## Data and materials availability

The code used for the simulations is available as a GitHub repository (https://github.com/MarcoColnaghi1990/LGT-repeat-sequences).

## Supplementary Information Text

### Supplementary Methods

We use a Fisher-Wright process with non-overlapping generations to model the evolution of a population of *N* haploid individuals, subjected to a deleterious point mutation rate of *μ* per locus per generation and undergoing transformation (LGT) at a rate *λ* (**Figure S1**). At the beginning of each simulation, every individual possesses a genome composed of *g* unique protein-coding genes, each of which can exist in either a wildtype or deleterious mutant state. The genome is assumed to be circular (locus *g* is contiguous with locus 1). The genome is interspersed at random intervals with a generic repeat sequence, at an initial density *ρ* per protein-coding gene. The initial positions of the repeats are randomly sampled from a uniform distribution (i.e., all loci regions have the same probability of harbouring a repeat sequence at the beginning of the simulation). For simplicity, protein-coding genes and repeats are treated as unitary entities, and we neglect sequence variability within these genes.

The new generation is obtained by sampling *N* individuals, with replacement, from the old population (**Figure S1**). The probability of reproduction is proportional to the individual fitness. Following previous theoretical studies (1-4), we assume no epistatic interactions and measure fitness as a multiplicative function

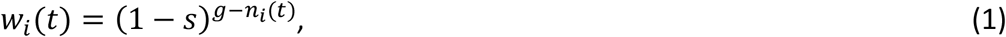

where *n*_*i*_*(t)* is the number of different functional protein-coding genes possessed by an individual *i* at time *t* (*g* − *n*_*i*_*(t)* is the number of genes that have been lost, either because of mutations or deletions). To avoid unnecessary complexity, we assume that gene duplication has a negligible effect on fitness, and only consider whether there is at least one functional copy of each protein-coding gene. Once a new generation is formed, the old generation dies, and their DNA forms the genetic pool from which the new generation acquires environmental DNA (eDNA) for recombination. We assume that eDNA strands are only stable for one generation before decaying irreversibly. Individuals of the new generation undergo recombination via LGT with a probability *λ*.

For each individual that undergoes LGT, a sequence of eDNA of length *L* is randomly sampled from the eDNA pool. The requisite for successful recombination is homology between the terminal loci of the eDNA sequence and the host genome. Homology can be either to a protein-coding gene or a repeat sequence. These dynamics follow experimental evidence that recombination of nonhomologous DNA can take place fairly easily in the presence of homologous flanking sequences, but not in their absence (5, 6); integration of homologous DNA is estimated to be at least a billion times more likely of that of strongly divergent “foreign” DNA (6).

After a sequence is sampled from the eDNA pool, one of its terminal loci is randomly selected and matched with a homologous locus *x*\* in the host genome. If multiple homologous loci are present, one site is selected at random. Homologous loci *x*_*k*_ to the other terminal locus of the eDNA are found in the recipient genome. For each *x*_*k*_ a recombination probability is generated according to a Gaussian distribution,

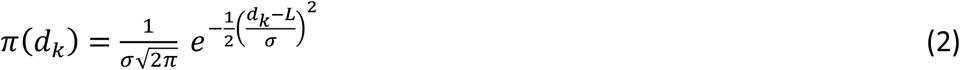

where *d*_*k*_ is the distance between *x*\* and *x*_*k*_, and the standard deviation is 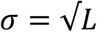 (where *L* is the length of the eDNA). This function reflects the assumption that longer eDNA sequences have greater variance in the distribution of recombination probabilities and penalises the probability of exchange over large distances (e.g., it is more likely that an eDNA sequence spanning 2 loci would recombine over a length spanning approximately 2 loci than approximately 20). This is in agreement with experimental data on recombination length in bacteria, showing the rarity of large recombination events and deletions (7, 8). Using Eq(2), one of the *x*_*k*_ loci is selected to be the other terminal region of recombination given weights proportional to *π(d*_*k*_). With successful pairing, all the elements between *x*\* and *x*_*k*_ (included) are substituted by the recombining eDNA sequence. If there is no match to either *x*\* or *x*_*k*_, recombination with the eDNA sequence is not possible and no genetic exchange takes place.

To model fully homologous recombination, we select two homologous loci *x*\* and *x*_*k*_ as above. If the eDNA and genomic sequences contain exactly the same genes in the same order (either as wild-type or mutant alleles) recombination successfully takes place (we assume that point mutations do not significantly affect the probability of homologous recombination). The sequence between *x*\* and *x*_*k*_ is excised and replaced by the eDNA sequence. Otherwise, no genetic exchange takes place.

After LGT, each individual acquires *m* new deleterious mutations, where *m* is a random integer drawn from a Poisson distribution with mean *U*. The genome wide mutation rate is given by *U* = *μg′*, where *g′* is the number of wildtype protein-coding genes, as we assume that mutated genes cannot mutate again. The position of the particular locus or loci in the genome that mutates is then randomly determined. For simplicity, the possibility of back mutation in protein-coding genes is neglected.

The evolutionary process is studied with a population initially free of mutants, over *t*_*max*_ = 5,000 generations, with 100 independent iterations for a given set of parameter values. For each replicate, we evaluate the gene-loss rate per generation from deletions (Δ*M*_*d*_/Δ*t)* and mutations (Δ*M*_*m*_/Δ*t)* as the average load of deletions and mutations respectively, divided by the number of generations. The total gene loss rate per generation Δ*M*/Δ*t* is calculated as the sum of these two components.

## Supplementary Figures

**Figure S1.**
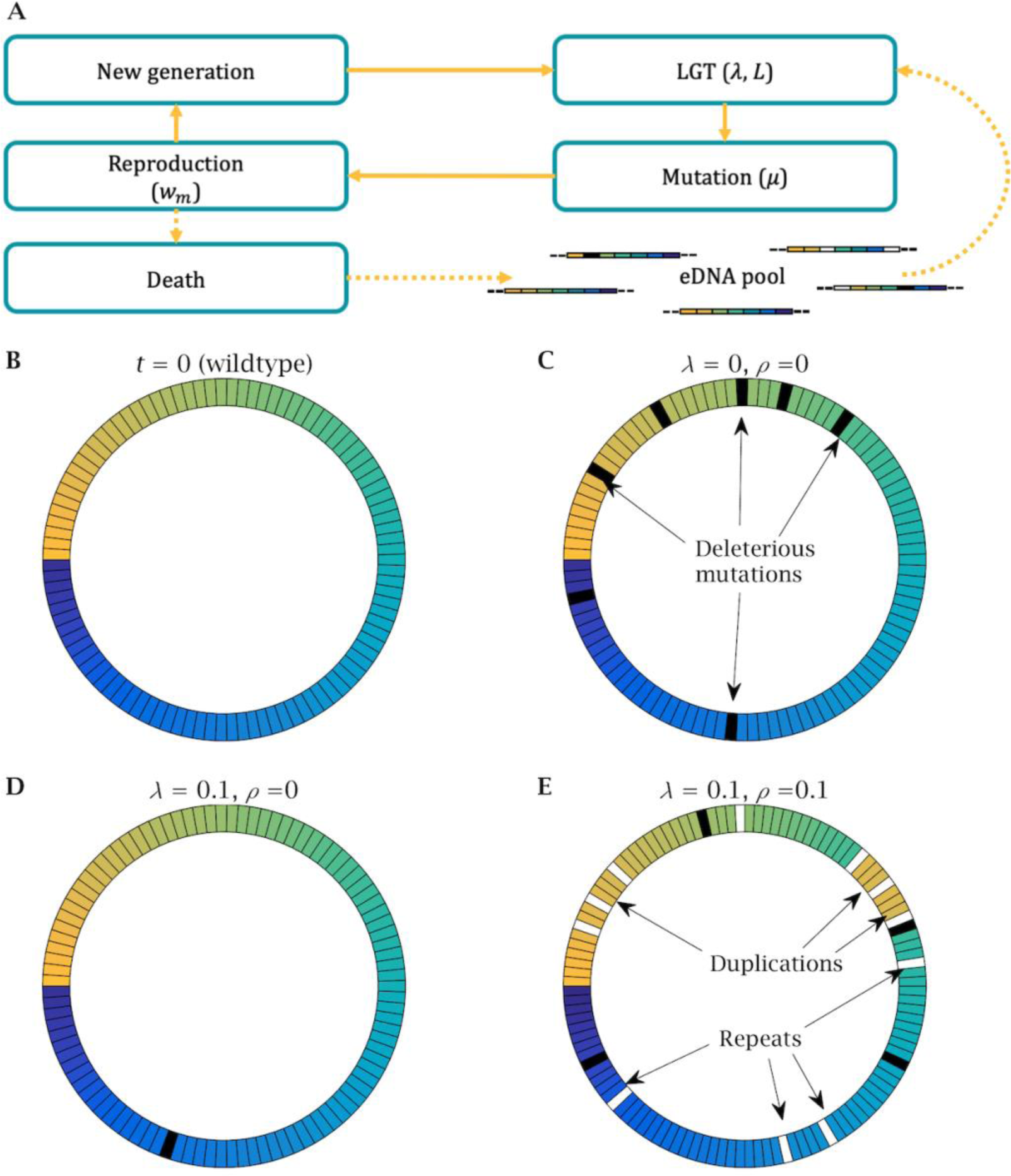
Illustration of the model dynamics. **(A)** Each generation, an individual has a probability *λ* of acquiring a fragment of eDNA of length *L* from the environment and recombining it. Following LGT, mutations are randomly introduced at a rate *μ* per locus. The new generation is then generated by sampling with replacement from the old generation in proportion to individual reproductive fitness *w*_*m*_. The old generation dies and its DNA is released into the environment and constitutes the eDNA pool for the new generation. **(B)** The wildtype genome is represented as a circular series of genes indicated by different colours. **(C)** The wildtype genome is subject to mutation pressure, resulting in the accumulation of deleterious alleles through Muller’s ratchet. **(D)** In a repeats-free population (*ρ* = 0), LGT (*λ* = 0.1) allows homologous recombination, increasing genetic variation and favouring the elimination of deleterious mutations. **(E)** In the presence of repeats (*ρ* = 0.1), the possibility of ectopic recombination limits the benefits of LGT, causing gene deletions and duplications. Other simulation parameters: *t* = 5,000, *g* = 100, *N* = 2,500, *L* = 5, *U* = 0.003.

**Figure S2.**
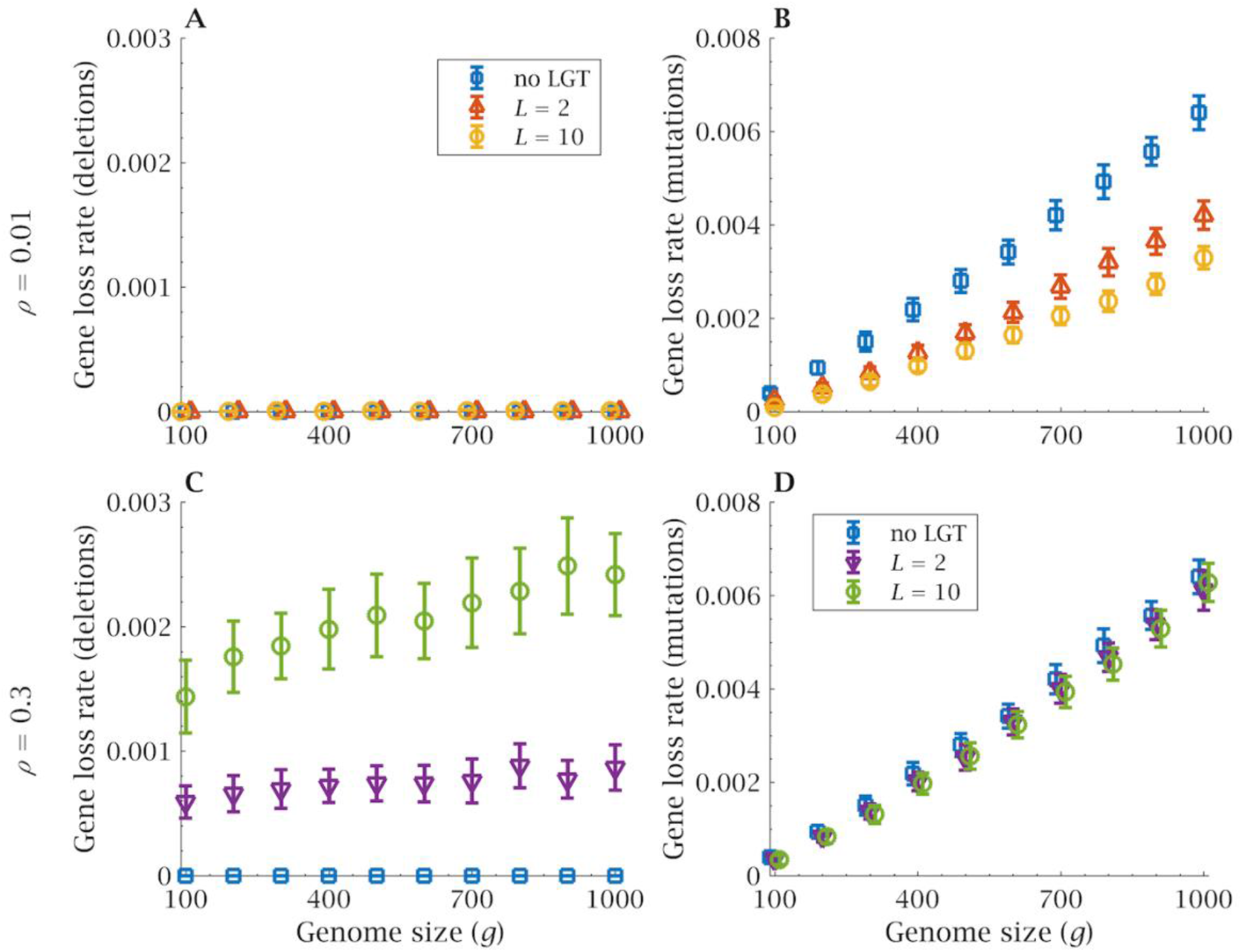
Genome size and recombination length. At low repeat density (*ρ* = 0.01), (**A**) increases in recombination length (*L*) limit mutation accumulation as genome size increase, (**B**) without increasing the rate of deletion. But in repeat-rich genomes (*ρ* = 0.3), (**C**) higher *L* provides virtually no benefits, (**D**) while introducing a large number of new deletions. Error bars show the standard deviation over 100 independent simulations. Gene loss rate was calculated over *t*_*max*_ = 5,000 generations. Other parameters: *N* = 2,500, *μ* = 10^−5^ and *λ* = 0.1.

